# Deep proteomic analysis of chicken erythropoiesis

**DOI:** 10.1101/289728

**Authors:** Marjorie Leduc, Emilie-Fleur Gautier, Anissa Guillemin, Cédric Broussard, Virginie Salnot, Catherine Lacombe, Olivier Gandrillon, François Guillonneau, Patrick Mayeux

## Abstract

In contrast to mammalian erythroid cells that lost their nucleus at the end of the differentiation process, circulating chicken erythrocytes, like erythrocytes of most other non-mammalian vertebrates, are nucleated although their nucleus is believed to be transcriptionally silent. This major difference suggests that the erythroid differentiation process is likely to present both similarities and differences in mammals compared to other vertebrates. Since proteins are the major cellular effectors, analysis of the proteome is more prone to reflect true differences than analysis of the pattern of mRNA expression. We have previously reported the evolution of the proteome of human erythroid cells throughout their differentiation process. Here we report the analysis of the proteome of chicken erythroblasts during their terminal differentiation. We used the T2EC cellular model that allows to obtain homogenous populations of immature erythroblasts. Induction of their terminal differentiation led to their maturation and the possibility to obtain cells at different differentiation stages. Mass spectrometry analysis of these cell populations allowed the absolute quantification of 6167 proteins throughout the terminal differentiation process. Beside many proteins with similar expression patterns between chicken and human erythroblasts, like SLC4A1 (Band3), GATA1 or CD44, this analysis also revealed that other important proteins like Kit or other GATA transcription factors exhibit fully different patterns of expression.

---

Although all vertebrates possess erythrocytes to facilitate oxygen and carbonic dioxide transports, their structures and physiology could be significantly different (Svoboda and Bartunek, 2015). Especially, mammalian erythroblasts expel their nucleus at the end of the differentiation process while circulating erythrocytes of almost all other vertebrates are nucleated. This is the case of the chicken erythrocytes that also contain mitochondria. Moreover, these cells present interesting specificities such as the expression of an erythroid-specific form of histone linker (H5) that is likely to play a role in the strong chromatin compaction that occurs during the differentiation of these cells (Kowalski and Palyga, 2016). In contrast to murine and human erythrocytes, chicken erythrocytes are believed to be devoid of glucose transporter, suggesting that they use other source(s) of metabolizable carbon (Johnstone et al., 1998). The control of erythrocyte production in chicken remains poorly understood. Chicken erythropoietin (Epo) has not been properly characterized and while a putative Epo receptor (EpoR) has been recently described (Hron et al., 2015), neither its biological function nor its pattern of expression has yet been identified.

Because of these specificities, the avian model has frequently been used to study specific aspects of erythropoiesis and robust biological models have been developed for this purpose (Beug et al., 1982; Gandrillon et al., 1999). High numbers of very immature erythroid progenitor cells can be obtained by culturing chicken bone marrow cells in the presence of TGF-α, TGF-β and dexamethasone. These T2EC cells can be maintained for up to 30 days in proliferation without differentiation. At any time, they can be induced to terminally differentiate into fully mature erythrocytes by addition of anemic chicken serum and withdrawal of TGF-α and TGF-β (Gandrillon and Samarut, 1998; Gandrillon et al., 1999).

The transcriptomes of T2EC cells in self renewing conditions and after 24h of differentiation have been previously compared by SAGE (Damiola et al., 2004) but no global analysis of the whole chicken erythroid differentiation process is currently available at the proteome level. Moreover, it is now clearly established that the relationship between the transcriptome and the proteome of a cell is rather low (see e.g. Schwanhausser et al., 2011). Indeed, variable mRNA translation efficiencies and protein stabilities partly disconnect mRNA and protein levels. In addition, it has been recently shown that a dynamic intron retention program is activated at the end of the mammalian erythroid differentiation process and we have recently shown that it could also participate to the discrepancy between mRNA and protein expression in erythroid cells (Gautier et al., 2016). Whether this intron retention process also occurs in chicken erythroblasts remains to be determined.

Current proteomic methods allow the quantification of thousand proteins starting from minute amount of cells (Kulak et al., 2017). We recently established the evolution of the proteome of human (Gautier et al., 2016) and murine (Gautier et al submitted) erythroid cells along the terminal differentiation process. A recent improvement of these methods allows the high throughput absolute quantification of proteins. This quantification method is based on the demonstration that the relative MS signal for a protein is roughly proportional to the relative mass abundance of this protein in the analyzed mixture (Wisniewski et al., 2014). The relative MS signal for a protein is defined as the sum of the MS intensities of all peptides attributed to this protein divided by the sum of the MS intensities of all identified peptides for all proteins. It has been subsequently demonstrated that histones provide an internal standard allowing the absolute quantification of cellular proteins with high accuracy. We have shown that this method allows the absolute quantification of erythroblast proteins with a good accuracy all along the terminal differentiation process.

In the present study, we have determined the evolution of the T2EC proteome during the terminal differentiation cells. Overall, we quantified 6167 proteins and their time-dependent evolution through 6 time points.

## Material and methods

### Cells

T2EC were grown and differentiated as previously described [1]. Briefly, cells were extracted from the bone marrow of 19 days-old SPAFAS white leghorn chicken embryos (INRA, Tours, France). They were grown in α–MEM medium supplemented with 10% foetal bovine serum (FBS), 1mM HEPES, 100 nM β-mercaptoethanol, 100 U/mL penicillin and streptomycin (PS), 5 ng/mL TGF-α, 1 ng/mL TGF-β and 1 µM dexamethasone. Differentiation was induced by placing cells into a differentiation medium made of α-MEM supplemented with 10% FBS, 1 mM HEPES, 100 nM β-mercaptoethanol, 100 U/mL PS, 10 ng/mL insulin and 5% anemic chicken serum (ACS; [2]). 2×10^6^ cells were sampled before differentiation induction (T0) and after 12h, 24h, 36h, 48h and 72h of differentiation, washed twice with PBS, flash frozen and sored at -80°C before processing.

### Sample preparation and mass spectrometry analysis

Sample preparation was done as previously described using the FASP procedure (Gautier et al., 2016). Briefly, cells were solubilized in 100µl of Tris/HCl 100 mM pH8.5 buffer containing 2% SDS. Total protein amounts were quantified using BCA (Pierce). 10 mM TCEP and 40 mM chloroacetamide were then added and the samples were boiled for 5min. 50µg of proteins were sampled and treated with urea to remove SDS. After urea removal, proteins were digested overnight with 1µg of sequencing-grade modified trypsin (Promega) in 50 mM Tris/HCl pH8.5 buffer. Peptides were recovered by filtration, desalted on C_18_ reverse phase StageTips and dried. They were then separated in 5 fractions by strong cationic exchange (SCX) StageTips (Kulak et al., 2014) and analyzed by mass spectrometry using an Orbitrap Fusion mass spectrometer (Thermo Fisher Scientific). Peptides from each fraction were separated on a C_18_ reverse phase column (2 μm particle size, 100 Å pore size, 75 μm inner diameter, 25 cm length) with a 170 min gradient starting from 99% of solvent A containing 0.1% formic acid in milliQ-H_2_O and ending in 55% of solvent B containing 80% ACN and 0.085% formic acid in milliQ-H_2_O. The mass spectrometer acquired data throughout the elution process. The MS1 scans spanned from 350 to 1500 Th with 1.10^6^ AGC target, 60ms maximum ion injection time (MIIT) and resolution of 60 000. MS Spectra were recorded in profile mode. HCD fragmentations were performed from the most abundant ions in top speed mode for 3 seconds with a dynamic exclusion time of 30 s. Precursor selection window was set at 1.6Th. HCD Normalized Collision Energy (NCE) was set at 30% and MS/MS scan resolution was set at 30000 with AGC target 1.10^5^ within 60ms MIIT. The mass spectrometry data were analyzed using Maxquant version 1.5.3.30 (Cox et al., 2014). The database used was a concatenation of chicken sequences from the Uniprot-Swissprot database (Uniprot, release 2017-07) and the list of contaminant sequences from Maxquant. Cystein carbamidomethylation was set as constant modification and acetylation of protein N-terminus and oxidation of methionine were set as variable modifications. Second peptide search and the “match between runs” (MBR) options were allowed. False discovery rate (FDR) was kept below 1% on both peptides and proteins. Label-free protein quantification (LFQ) was done using both unique and razor peptides with at least 2 such peptides required for LFQ. Absolute quantifications were done using histones as standard (Wisniewski et al., 2014). Statistical analysis and data comparison were done using the Perseus software (Tyanova et al., 2016).

## Results

T2EC (TGF-α, TGF-β-induced erythrocytic cells) is a robust cellular model studied for several decades. Those are chicken erythroid progenitors derived from normal bone marrow cells extracted from the tibias of 19-day-old SPAFAS chicken embryos which grow in the presence of TGF-α, TGF-β and dexamethasone (Gandrillon et al., 1999). This specific combination of growth factors acts at the progenitor level and allows a stable growth of self-renewing cells for about 30 days before the cells reach their Hayflick limit and stop proliferating. The presence of TGF-β in the culture medium leads to a strong inhibition of the growth of non-erythroid adherent cells, like fibroblasts. In the self-renewal conditions, there is no overt differentiation (less than 0.1% of benzidine positive cells). The differentiation of these cells is induced by the concomitant withdrawal of growth promoting factors (TGF-α, TGF-β and dexamethasone) and the addition of anemic chicken serum (Gandrillon and Samarut, 1998) plus insulin. We determined the evolution of the proteome of these cells along the differentiation process through 6 time points from 0 to 72h.

Overall, we identified 6167 proteins with a FDR below 1% at the peptide and protein level (Table S1, sheet “Overall data”). We used the sum of the LFQ signals of histone peptides to calculate the absolute expression of these proteins. The histone protein mass per cell has been shown to be roughly similar to the mass of DNA (Wisniewski et al., 2014). We determined that T2EC cells contained 4.4 pg DNA per cell and we used this value to calculate the cellular mass of each identified protein per cell starting from the LFQ values of each protein. Absolute quantification was obtained for 6137 proteins (Table S1, sheet “Absolute quantifications”). The number of proteins quantified at each differentiation stage varied between from 4972 proteins (cells at 72h of differentiation) to 5415 proteins (cells at 12 hours of differentiation, figure 1A). 4475 proteins were quantified at each differentiation stage and 756 proteins were only quantified at a single differentiation stage (figure 1B). As expected since the size of the cells decreases during the differentiation process, the overall protein mass of fully differentiated cells sampled after 72h of differentiation induction was around two folds lower than the protein mass of undifferentiated cells (figure 1C). Accordingly, this is accompanied by a massive down-regulation of the majority of the proteins (figure 1D).

**Figure 1:**
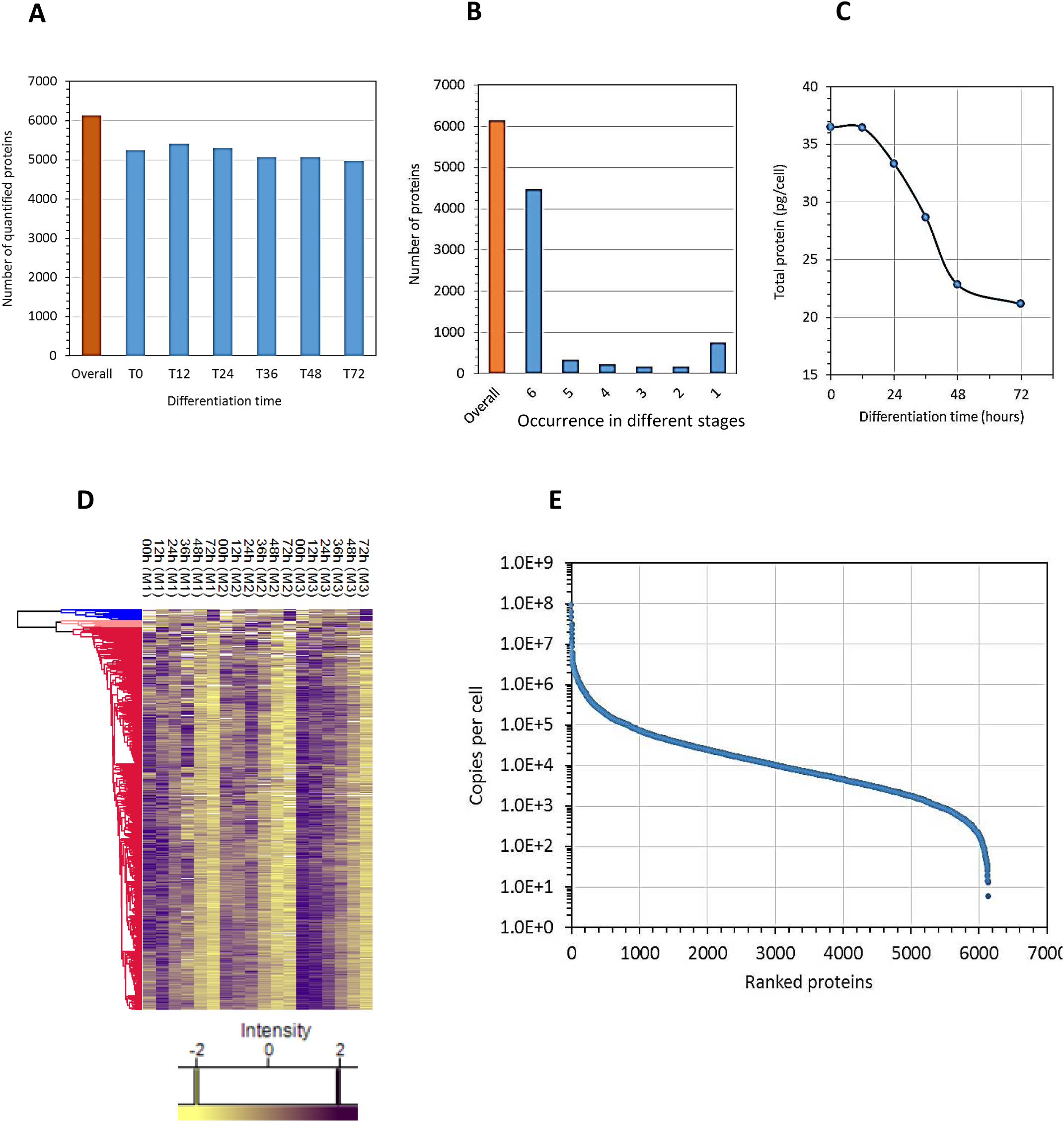
Global proteomic data analysis: 1A, number of quantified proteins at each differentiation stage, the brown bar indicate the total number of quantified proteins. 1B, number of quantified proteins according to the number of differentiation stages in which they have been quantified. 1C, total cellular amount of protein at each differentiation stage. The total amount of proteins as been determined from the absolute quantifications by summing the amounts of each quantified proteins. 1D: the gene were regrouped by hierarchical clustering, showing a small group of genes (in blue) the expression of which globally increases during the differentiation, a small group in pink witch increases and then decreases and a huge majority (in red) of genes the expression of which decreases during differentiation. 1E: dynamic range of protein expression, the quantified proteins were ranked according to their determined copy numbers.

The range of quantified proteins extended through seven logs from nearly 100 million copies per cell for histones and for globins at the end of the differentiation process up to proteins expressed at a few copies per cell (figure 1E). The patterns of expression of classical differentiation markers showed both similarity and discrepancies with their expression during mammalian erythropoiesis. As in mammalian CFU-E, globins are already expressed at the undifferentiated stage and their levels increased dramatically during differentiation as expected (figure 2A and B). Band3 (SCL4A1) expression also strongly increased as observed during murine or human erythropoiesis (figure 2C). CD 44 expression decreases, as observed during murine erythropoiesis especially (figure 2D). In sharp contrast the expression of Kit increases during the erythroid differentiation process in the chicken (figure 2E) while its expression strongly decreases during mammalian erythropoiesis. Indeed, Kit is no longer detectable after the basophilic erythroblast stage in murine or human erythroblasts. Interestingly, we similarly did not detect the putative EpoR that has been recently identified in the chicken genome (Hron et al., 2015). We also did not detect the expression of this putative EpoR protein in the 6C2 and HD3 chicken cell lines (data not shown).

**Figure 2:**
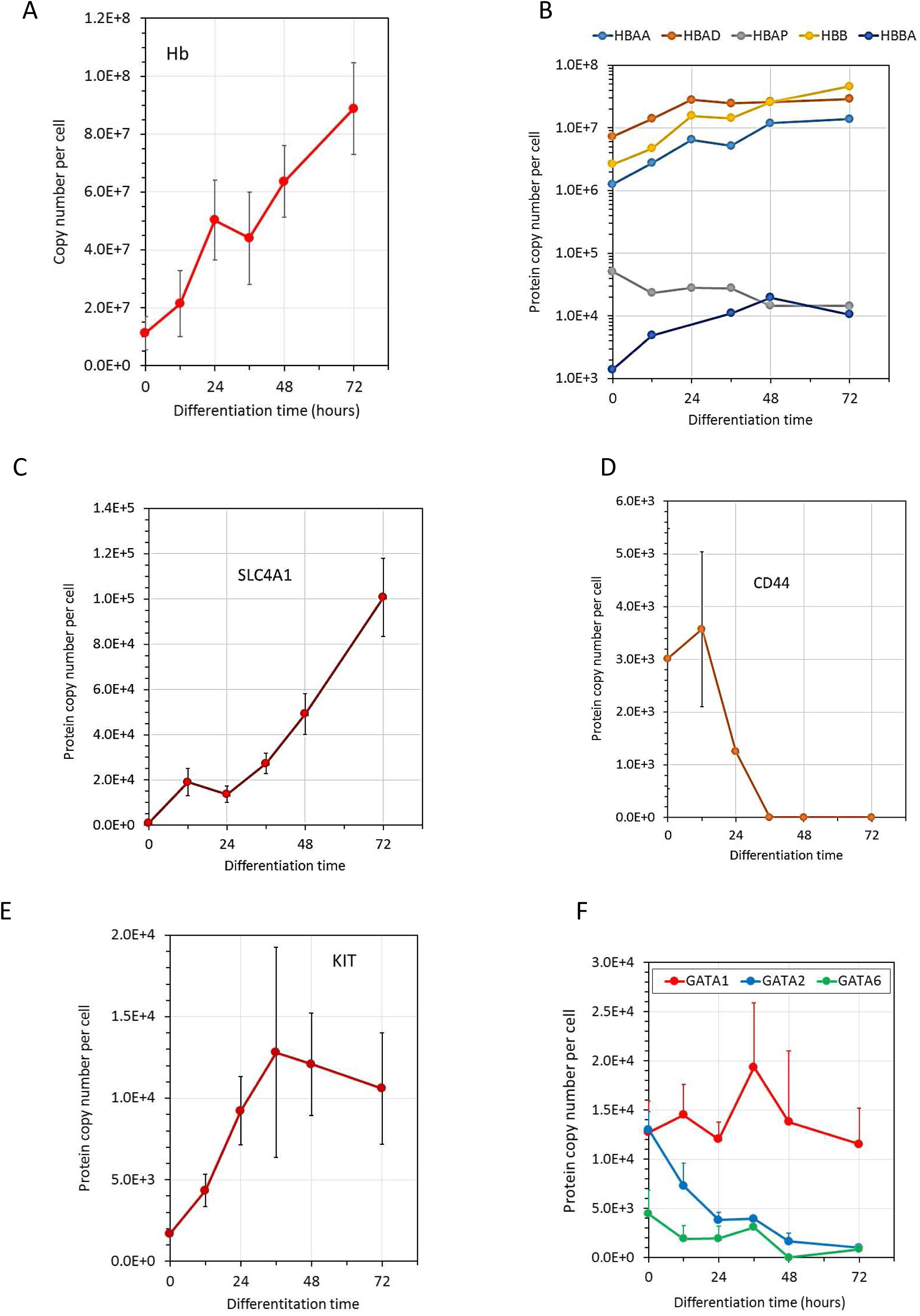
Specific expression of marker proteins: 2A, the number of copies of each globin chains was summed. 2B expression of individual globin chains in a logarithmic scale, 2C, D, E and F expression of Band3 (Slc4A1), CD44, Kit and GATA proteins, respectively.

Gata1 is expressed before the induction of terminal differentiation and its level did not significantly increase during the differentiation. Gata2 is expressed at the same level as Gata1 in undifferentiated cells and its expression decreases during terminal differentiation. Roughly similar patterns of GATA1 and GATA2 expression were observed during mammalian erythropoiesis, although GATA2 expression decreases more sharply and is completely abolished as soon as the cells enter the terminal differentiation process. Surprisingly, chicken erythroblasts express GATA6 that is not detected in mammalian erythroblasts (figure 1F).

Finally, we assessed the level of correspondence between mRNAs and proteins in our cells. For this we extracted the RNA mean expression value during T2EC differentiation from single cell transcriptomics (Richard et al., 2016). 48 genes were found in both the proteome and transcriptome dataset and their expression was compared. Among those genes, only 17 were found to be significantly correlated (figure 3A and 3B for some examples), showing the importance of complementing a transcriptome analysis with a proteomic analysis.

**Figure 3:**
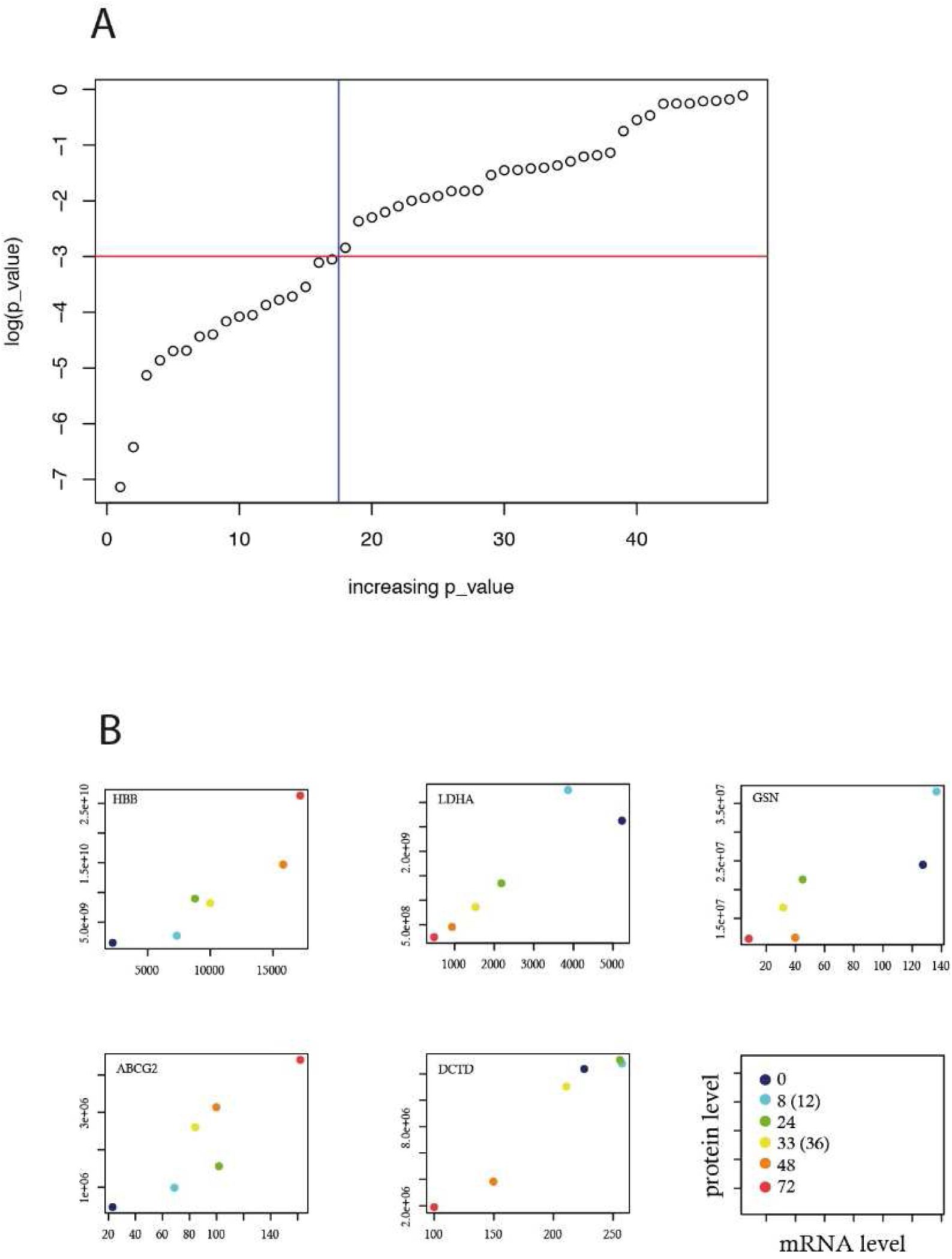
Quantitative comparison between transcriptome and proteome evolution during differentiation: 48 genes were found both in proteomic data (this study) and transcriptomic data (Richard et al., 2016). The significance of their Pearson correlation was assessed by the cor.test function using R (R_Core_Team, 2017). A: 17 genes were displaying a p-value of less the 0.1%. B: 5 genes significantly correlated are shown. The right bottom panel describes the value of the axis as well as the differentiation time points compared.

## Discussion

Our data provide the first description of the proteome evolution of erythroblasts during their differentiation in a non-mammalian species. For this analysis, we have used the T2EC model of erythroid differentiation. This model has been previously extensively characterized (Gandrillon et al., 1999). The erythroid nature of these cells was first ascertained by a very high expression level of the transferrin receptor, recognized by the JS8 antibody (Gandrillon et al., 1999), a result confirmed by the proteomic studies since these cells express more than one million of transferrin receptor molecules at the onset of terminal differentiation (table S1). The absence of mature myeloid or lymphoid cells in the cultures was ascertained by the absence of detectable expression of the transcription factor Spi-1/PU.1 by RT-qPCR (Gandrillon et al., 1999) as well as by the absence of specific antigens for the eosinophilic, myelomonocytic and thrombocytic lineages (Dazy et al., 2003). The differentiation is characterized by a strong reduction of cell size which affects all the cells, confirming the erythroid nature of those cells (Gandrillon et al., 1999). Our data show that, as expected, the total amount of cellular proteins simultaneously drops (figure 1C). Benzidine staining, allowing to measure the percent of hemoglobin-expressing cells, indicates a rapid increase of mature cells which reaches up to 100% at 6 days after induction of differentiation (Gandrillon et al., 1999). Altogether, these data demonstrate that the T2EC model mostly contains erythroid precursors, whose differentiation is blocked at the ProE stage, with few if any contamination with non-erythroid cells. The differentiation of these cells is rather synchronous and cells differentiate up to the erythrocytic stage in culture.

Chicken erythropoiesis present several specificities. Indeed, despite extensive research, chicken Epo and EpoR genes are not definitively identified. The pattern of Kit expression in chicken erythroblasts is surprising and raises the question of the role of SCF, the Kit ligand, in avian terminal differentiation. Indeed, during mammalian erythropoiesis, Kit is mainly expressed during the amplification phase of erythropoiesis and SCF is required for an efficient cell proliferation. Its expression is rapidly down regulated after the ProE stage and Kit is no longer detectable after the basophilic stage while its expression strongly increases during the terminal differentiation of chicken erythroblasts. The clear identification of the avian EpoR is mandatory to analyze the cooperation between the two growth factors, Epo and SCF, and their respective roles in avian erythropoiesis.

## Author contributions

AG and OG realized the cell cultures and characterized the erythroid cells, ML, EFG, CB and VS realized the mass spectrometry analyses, ML, EFG, CB, VS, OG, FG and PM analyzed the proteomic data. FG, OG and PM managed the study, PM and CL wrote the article, all authors discussed data and critically reviewed the manuscript.

**Supplementary table S1:** Overall proteomic data (Excel file)

